## Supplementary figures and images for "TTLL12 is required for primary ciliary axoneme formation in polarized epithelial cells"

### Supplemental Figure 1

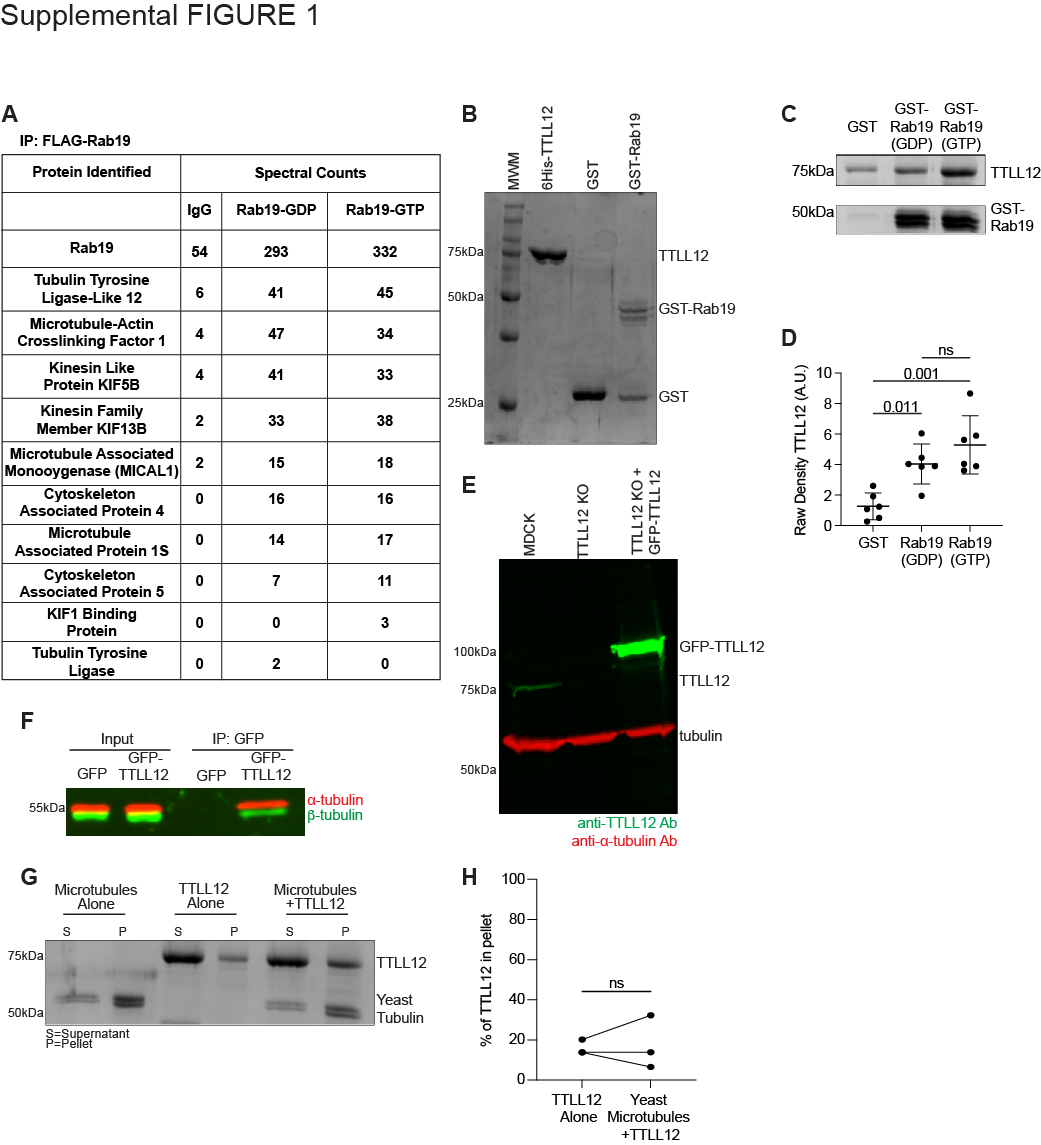

### Supplemental Figure 2

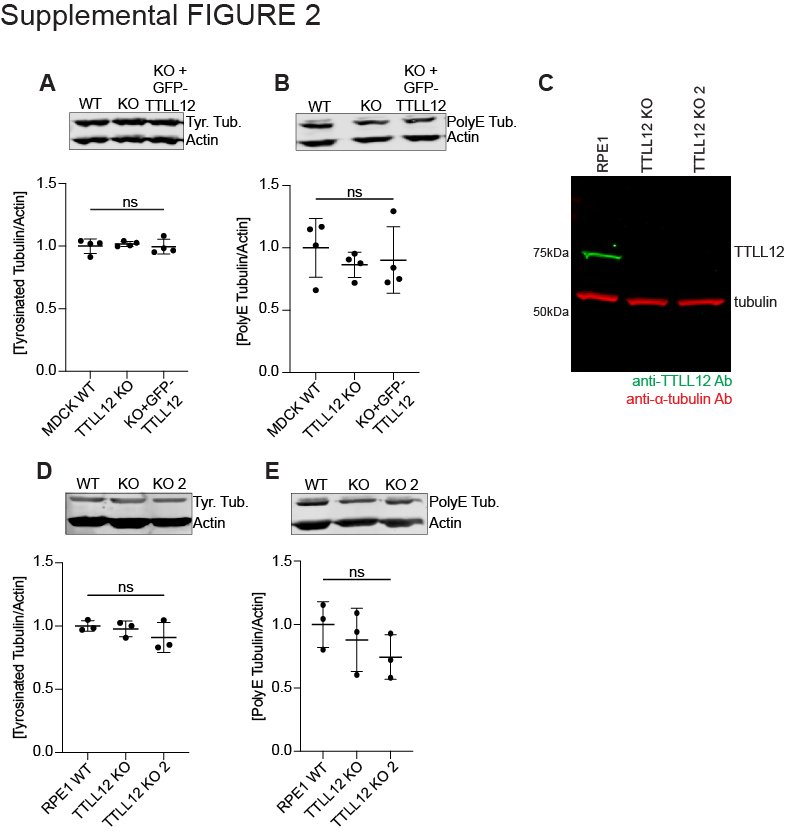

### Supplemental Figure 3

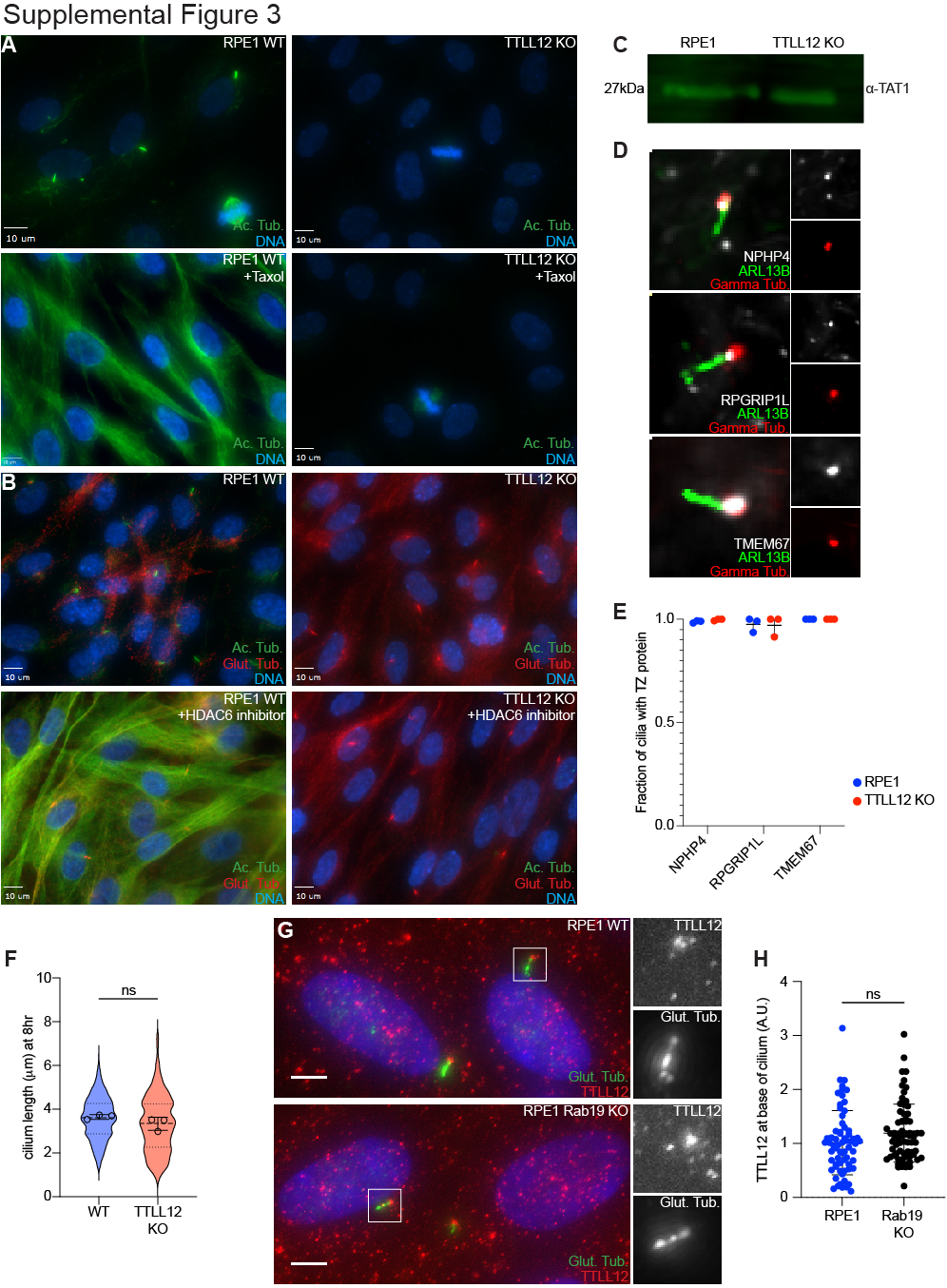

### Supplemental Figure 4

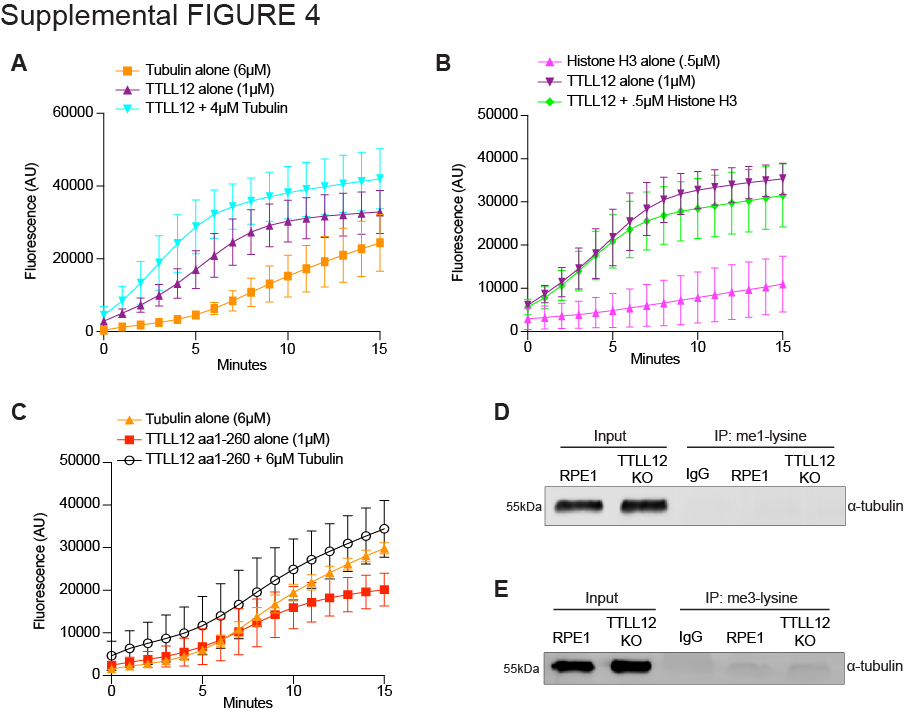
